# Preliminary Insights into the Acute Molecular Responses in C2C12 Myotubes to Hyperthermia and Insulin Treatment

**DOI:** 10.1101/2025.03.26.644592

**Authors:** Ross Gillette, Irene C. Turnbull, Venugopalan D. Nair, Angelo Gaitas

**Affiliations:** Department of Neurology, Icahn School of Medicine at Mount Sinai, New York, NY, USA; Cardiovascular Research Institute, Icahn School of Medicine at Mount Sinai, New York, NY, USA; BioMedical Engineering & Imaging Institute, Icahn School of Medicine at Mount Sinai, New York, NY, USA

**Author notes:** These authors contributed equally to this work (Ross Gillette, Irene C. Turnbull).

**Keywords:** hyperthermia, insulin, myotubes, skeletal muscle, temperature, stress

## Abstract

This study investigates the differential gene expression in an immortalized cell line of mouse skeletal myoblasts (C2C12)-derived myotube cells subjected to hyperthermia (40°C) with and without insulin treatment to elucidate the impact of thermal stress on skeletal muscle physiology. Hyperthermia, which occurs during intense physical activity or environmental heat exposure, is known to challenge muscle homeostasis and influence metabolic function. mRNA sequencing revealed that hyperthermia robustly altered gene expression—upregulating key genes involved in glycolysis, oxidative phosphorylation, heat shock response, and apoptosis. These changes are suggestive of an elevated metabolic state and enhanced cellular stress; however, these results remain preliminary without complementary protein or metabolic assays. Notably, insulin treatment moderated many of the hyperthermia-induced transcriptional alterations, particularly affecting genes linked to glucose uptake and metabolism. Together, these findings provide hypothesis-generating insights into the interplay between thermal stress and insulin signaling in C2C12 myotubes, and they underscore potential targets for future mechanistic studies.

## 1. Introduction

Maintaining cellular homeostasis is essential for proper physiological function, yet cells are continually challenged by stressors such as hyperthermia and hormonal fluctuations [1,2]. Hyperthermia—whether due to fever or environmental heat stress—triggers adaptive responses including the induction of heat shock proteins, metabolic reprogramming, and apoptosis [3-5]. Insulin, a key regulator of glucose metabolism, modulates glucose uptake and energy homeostasis. Although the individual effects of hyperthermia and insulin have been studied, their combined impact on cellular processes in skeletal muscle remains poorly understood [6-9].

C2C12 skeletal myoblasts, under prescribed conditions, differentiate and fuse into multinucleated myotubes, and serve as in vitro models of skeletal muscle cells [10]; C2C12 myotubes are particularly sensitive to both thermal and hormonal signals [11-14]. This sensitivity makes them an ideal model for investigating how hyperthermia and insulin interact to influence cellular function. Importantly, understanding this interplay may be relevant to metabolic disorders such as diabetes, where dysregulated glucose homeostasis plays a central role [15-23].

In this study, we employed mRNA sequencing (RNA-seq) to evaluate the acute molecular responses of C2C12 myotubes subjected to hyperthermia (40°C) with and without insulin. By identifying key gene networks and pathways modulated under these conditions, our results generate testable hypotheses regarding the molecular mechanisms that govern energy metabolism and stress responses in C2C12 cells. While our findings highlight intriguing transcriptional shifts, they also underscore the need for future studies incorporating protein-level and functional assays to fully elucidate the cellular outcomes of combined thermal and insulin stress.

## 2. Materials and Methods

### 2.1 Cell Culture

C2C12 myoblast cells were obtained from the American Type Culture Collection (ATCC) and cultured in Dulbecco’s Modified Eagle Medium (DMEM) with 4.5g/l glucose (Corning Cat # 10-013-CV) supplemented with 10% fetal bovine serum (FBS) (Corning, Cat#:35-011-CV) and 1% penicillin-streptomycin (Corning, Cat# 30-002-CI). Cells were maintained at 37°C in a humidified atmosphere with 5% CO2. For differentiation, cells were seeded in 6-well plates and grown to ∼80% confluence before switching to differentiation medium (DMEM with 4.5 g/L glucose, 2% horse serum [Corning, Cat #:35-030-CV], and 1% penicillin-streptomycin). The medium was refreshed every other day until day 8, when myotube formation was confirmed (Fig. 1).

**Figure 1.**
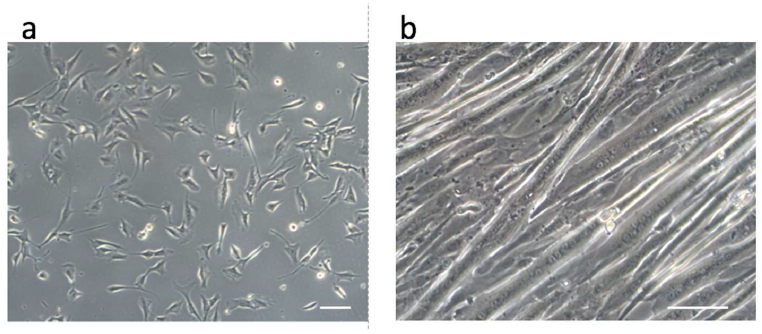
Myoblasts and myotubes in culture. Representative microphotographs of myoblasts (**a**), and myotubes (**b**). Scale bar 100 micron.

### 2.2 Hyperthermia and Insulin Treatment

To synchronize the cell cycle and reduce basal metabolic activity, cells were serum-starved overnight in DMEM containing 0.5% FBS. Subsequently, cells were divided into four groups (n = 3 per group) as follows. 1) Control: maintained at 37°C, 2) Hyperthermia: exposed to 40°C for 30 minutes, 3) Insulin: treated with 100 nM insulin at 37°C for 30 minutes (Millipore Sigma, Cat #I9278), 4) Combined treatment: exposed to 40°C with 100 nM insulin for 30 minutes. (Fig. 2). Hyperthermia was achieved by moving the plated cells from the 37°C incubator and into an incubator with temperature set at 40°C with 5%CO2 for the prescribed time (30 minutes). The insulin treatment was applied (to the indicated groups) immediately before placing the cells either back into the 37°C incubator or into the 40°C incubator (hyperthermia). Immediately following treatment, cells were harvested for RNA extraction.

**Figure 2.**
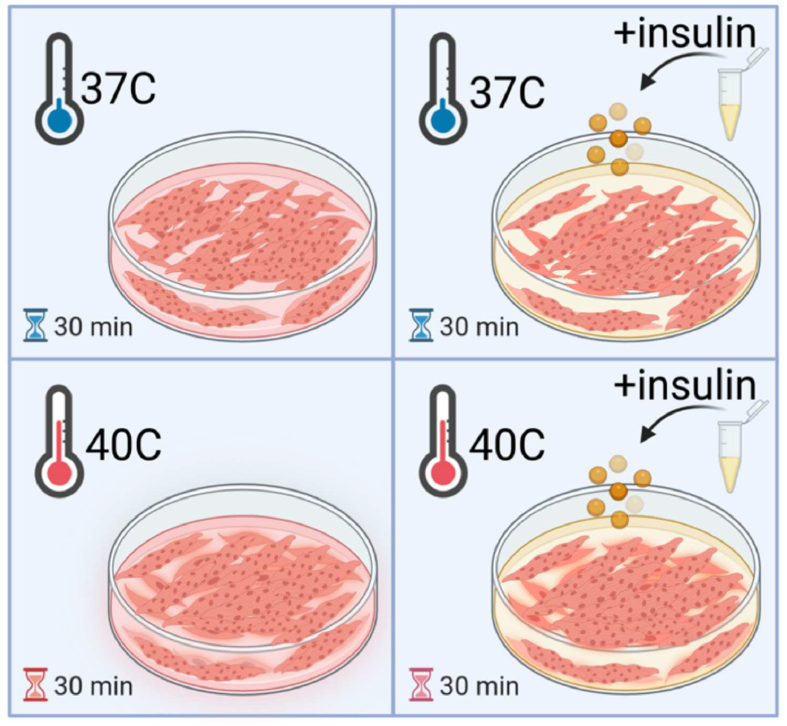
Experimental Design. Schematic of the four different conditions that were tested.

### 2.3 RNA Extraction and Sequencing

Total RNA was extracted using Direct-zol RNA extraction kit with Tri-reagent (Zymo Research, Cat# R2051) following the reported protocol [24]. RNA quality and concentration were assessed using the Agilent 2100 Bioanalyzer and NanoDrop spectrophotometer. RNA samples from one independent experiment with an RNA integrity number (RIN) above 7.0 were used for library preparation.

RNA-seq libraries were prepared using the Universal Plus mRNA-Seq kit (NuGEN/Tecan), according to the manufacturer’s protocol. Briefly, mRNA was isolated from total RNA, fragmented, and reverse transcribed to cDNA. Adapters were ligated, and the cDNA was amplified by PCR. The resulting libraries were quantified, pooled, and sequenced on an Illumina NextSeq 2000 platform, targeting an average of 35 million read pairs (70 million paired-end reads) per sample using a paired-end 100 base pair run configuration.

### 2.4 Bioinformatics and Differential Expression Analysis

#### 2.4.1 Quality Control and Alignment

Raw sequencing reads were subjected to quality control using FastQC to assess read quality, adapter content, and duplication rates. Low-quality bases and adapter sequences were trimmed using Trim_galore (v4.6) [25], a wrapper for Cutadapt (v4.6) [26]. Reads shorter than 20 nt or with a quality score <20 were excluded from further analysis. The trimmed reads were then aligned to the Mus musculus reference genome (mm10, GRCm38) using TopHat (v2.1.1) [27] and bowtie2 (v2.5.1) [28]. To reduce PCR bias, unique molecular identifiers (UMIs) were appended to the aligned reads using fgbio (v2.2.0) [29] and deduplicated using umi_tools (v1.1.4) [30]. For paired-end data, adapters were also trimmed using Trim_galore [25] a convenience wrapper for Cutadapt (v4.6) [26]. Additionally, high-quality reads were aligned using STAR aligner.

#### 2.4.2 Gene Count Normalization

Gene counts were generated using the HTSeq.scripts.count (v1.05) [31] in Python (v3.10.13) [32]. Contaminant reads were identified by aligning to the PhiX and hg38 (GRCh38) genomes, as well as to ribosomal and mitochondrial RNA sequences from mm10, using bowtie2 (v2.5.1) [28]. File formatting and sorting for downstream analyses were performed with samtools (v1.18) [33]. Quality control reports were generated at each step using FastQC (v0.12.1) followed by MultiQC (v1.18) [34]. Low expressing genes were removed early in analysis and before normalization using the filterByExpr function in the edgeR package.

Normalization and differential expression analysis were performed in R (v4.3.2) using the edgeR package (v4.0.12) [35] with the Trimmed Mean of M values (TMM) method [36, 37]. Biological replicates were grouped by identity and the desired contrasts (discussed below), and statistical model were established with the makeContrasts function in edgeR. Dispersion estimates were then recalculated, a quasi-likelihood generalized linear model [38] was fitted, and finally, statistical testing was performed with the contrasts and model previously mentioned.

#### 2.4.3 Statistical Analysis

A general lineal model was used to determine differential expression across a treatment type. These group-wise statistical comparisons were defined as follows:

Primary comparisons:

- Insulin effect: 37°C vs. 37°C + Ins.
- Temperature effect: 37°C vs. 40°C.
- Combined treatment effect: 37°C vs. 40°C + Ins.

Additional comparisons evaluated the temperature effect in the context of insulin (37°C + Ins vs. 40°C + Ins) and the insulin effect at hyperthermia (40°C vs. 40°C + Ins).

Differentially expressed genes (DEGs) were called using the decideTests() function in limma package for R. Multiple comparisons were corrected for using the Benjamini-Hochberg false discovery rate (FDR) method. An FDR cut off was set at FDR < 0.05.

### 2.5 Pathway and Gene Ontology Analysis

Gene Ontology (GO) and KEGG pathway enrichment analyses were performed in R on the DEGs [37,39]. Enrichment was considered significant at an FDR < 0.05.

## 3. Results

### 3.1. Quality Control

Sequencing libraries exhibited high quality, with Phred scores consistently above 30, indicating high base call accuracy. Of the three biological samples per group, one biological sample from the control group (which showed significant PCR duplication rates during deduplication) was excluded from further analysis to avoid skewing normalization and statistical power. The remaining samples demonstrated satisfactory alignment rates to the Mus musculus reference genome (mm10), ranging from 85% to 95%, with minimal off target alignments that were discarded.

### 3.2. Differential Gene Expression Analysis

Principal component analysis (PCA) identified two primary components that account for 50.48% and 13.01%, respectively (Supplementary fig. S1a). Pair-wise plotting of sample scores revealed that the first component predominantly separated the 40°C samples, which clustered tightly together, whereas the control (37°C) samples and those receiving insulin treatment were more dispersed (Supplementary fig. S1b).

Transcriptome-wide analysis showed distinct expression trends across primary comparisons (Fig. 3a); notably, hyperthermia alone resulted in a more dispersed pattern with an increased number of downregulated genes (Fig. 3b).

**Figure 3.**
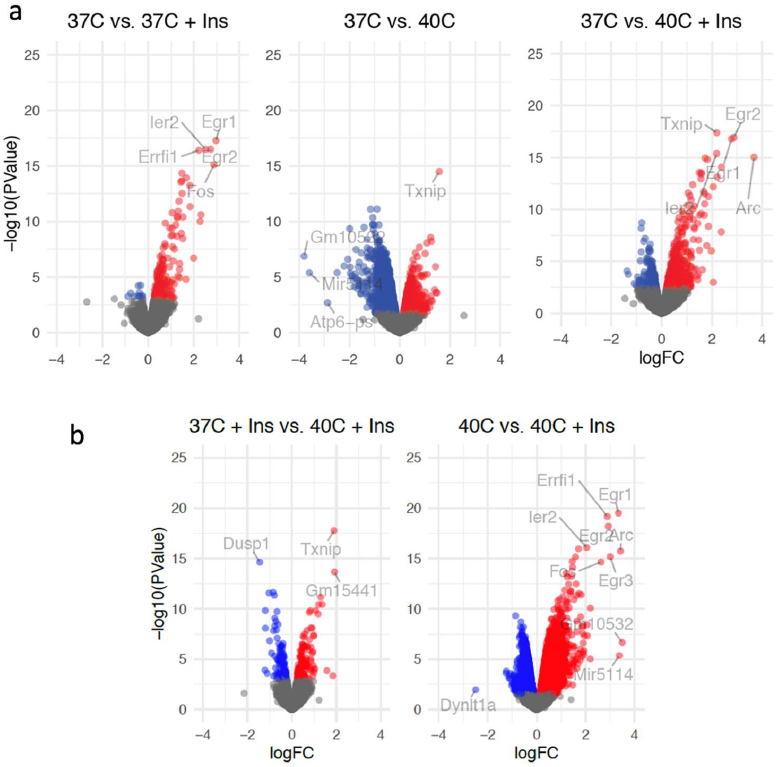
Volcano plots showing the differentially expressed genes between different comparisons. **a**) Volcano plots show transcriptome wide differentially expressed genes between control (37°C) and hyperthermia (40°C, without and with insulin). **b**) Volcano plots show transcriptome wide differentially expressed genes in balancing comparisons. Blue indicates downregulated genes, red indicates upregulated genes. Biological replicates: n=2 for 37°C group, and n=3 for all other groups.

Hyperthermia (40°C) induced widespread transcriptional changes, with 1772 genes upregulated and 2517 downregulated relative to the control (37°C) (Table 1). Many altered genes are involved in glucose metabolism, stress responses, and apoptosis. For example, downregulation of *Errfi1, Fos*, and *Egr*, suggests reduced proliferative and adaptive signaling under thermal stress (Table 2), whereas upregulation of *Txnip* indicates a shift toward managing oxidative stress and modulating glucose utilization (Table 2). Additionally, glycolytic enzymes such as *Gapdh, Tpi1*, and *Pfk1* were differentially expressed (Table 3), and several electron transport chain components (e.g., *Ndufa1, Cox6a1, Atp5mc3, Atp5mk, Atp5pb, Atp5mg, Atp5f1b, Atp5f1d*, and *Atp5f1e*) were upregulated (Supplemental data table S1). Hyperthermia also significantly increased expression of reactive oxygen species (ROS) response genes (*Sod1, Gpx1, Hmox1*) and apoptotic genes (Bok, Trp53, Bbc3) (Supplemental data table S1).

**Table 1.** Summary of number of DEGs.

|  | 37+ Ins | 40C | 40C+ Ins | 40C+ Ins | 40C+ Ins |
| --- | --- | --- | --- | --- | --- |
|  | 37C | 37C | 37C | 37C+ Ins | 40C |
| Up | 193 | 1772 | 694 | 136 | 3795 |
| N.S. | 12191 | 8106 | 11419 | 12156 | 5932 |
| Down | 11 | 2517 | 282 | 103 | 2668 |

Note: N.S. = not significant. Number of DEGs identified for each group-wise statistical comparison.

**Table 2.** Largest magnitude DEGs.

|  | 37+Ins |  | 40C |  | 40C+Ins |  | 40C+Ins |  | 40C+Ins |  |
| --- | --- | --- | --- | --- | --- | --- | --- | --- | --- | --- |
|  | 37C |  | 37C |  | 37C |  | 37C+Ins |  | 40C |  |
|  | logFC | FDR | logFC | FDR | logFC | FDR | logFC | FDR | logFC | FDR |
| Arc | 2.31 | **0.0000\*** | 0.25 | 0.4140 | 3.67 | **0.0000\*** | 1.36 | **0.0000\*** | 3.43 | **0.0000\*** |
| Dusp1 | 0.66 | **0.0000\*** | -0.65 | **0.0000\*** | -0.78 | **0.0000\*** | -1.44 | **0.0000\*** | -0.14 | 0.1208 |
| Dynlt1a | 2.20 | 0.4080 | 2.55 | 0.0729 | 0.05 | 0.9855 | -2.15 | 0.3285 | -2.49 | **0.0248\*** |
| Egr1 | 2.97 | **0.0000\*** | -0.57 | **0.0002\*** | 2.78 | **0.0000\*** | -0.19 | 0.3285 | 3.34 | **0.0000\*** |
| Egr2 | 2.73 | **0.0000\*** | -0.06 | 0.7184 | 2.87 | **0.0000\*** | 0.14 | 0.4984 | 2.93 | **0.0000\*** |
| Egr3 | 1.15 | **0.0000\*** | -1.30 | **0.0000\*** | 1.71 | **0.0000\*** | 0.56 | **0.0037\*** | 3.01 | **0.0000\*** |
| Errfi1 | 2.22 | **0.0000\*** | -1.07 | **0.0000\*** | 1.82 | **0.0000\*** | -0.40 | **0.0011\*** | 2.89 | **0.0000\*** |
| Fos | 2.88 | **0.0000\*** | -0.58 | **0.0099\*** | 2.05 | **0.0000\*** | -0.83 | **0.0000\*** | 2.63 | **0.0000\*** |
| Gm10532 | -0.29 | 0.7474 | -3.80 | **0.0000\*** | -0.30 | 0.6124 | -0.01 | 0.9942 | 3.50 | **0.0000\*** |
| Gm15441 | 0.29 | 0.3729 | 1.26 | **0.0000\*** | 2.21 | **0.0000\*** | 1.91 | **0.0000\*** | 0.95 | **0.0000\*** |
| Ier2 | 2.51 | **0.0000\*** | 0.14 | 0.2545 | 2.19 | **0.0000\*** | -0.32 | **0.0193\*** | 2.05 | **0.0000\*** |
| Mir5114 | -0.17 | 0.8928 | -3.59 | **0.0002\*** | -0.22 | 0.7882 | -0.05 | 0.9778 | 3.38 | **0.0001\*** |
| Txnip | 0.30 | **0.0493\*** | 1.57 | **0.0000\*** | 2.19 | **0.0000\*** | 1.90 | **0.0000\*** | 0.62 | **0.0000\*** |

Note: FC: fold-change; FDR: false discovery rate. Values in bold(\*) indicate the genes selected as significant from the corresponding comparisons.

**Table 3.** Glycolysis pathway genes.

|  | 37+Ins |  | 40C |  | 40C+Ins |  | 40C+Ins |  | 40C+Ins |  |
| --- | --- | --- | --- | --- | --- | --- | --- | --- | --- | --- |
|  | 37C |  | 37C |  | 37C |  | 37C+Ins |  | 40C |  |
|  | logFC | FDR | logFC | FDR | logFC | FDR | logFC | FDR | logFC | FDR |
| Aldoa | -0.07 | 0.7323 | 0.20 | 0.0506 | -0.17 | 0.1941 | -0.09 | 0.6938 | -0.37 | **0.0003\*** |
| Eno1 | -0.06 | 0.7550 | 0.12 | 0.1790 | -0.08 | 0.4906 | -0.02 | 0.9413 | -0.20 | **0.0093\*** |
| Eno1b | -0.15 | 0.4038 | 0.06 | 0.5506 | -0.21 | 0.0767 | -0.06 | 0.8083 | -0.27 | **0.0019\*** |
| Eno2 | 0.05 | 0.8182 | -0.06 | 0.5546 | -0.03 | 0.8347 | -0.09 | 0.7206 | 0.03 | 0.7409 |
| Eno3 | -0.12 | 0.5872 | 0.19 | 0.0707 | -0.23 | 0.0872 | -0.11 | 0.6205 | -0.42 | **0.0001\*** |
| Gapdh | -0.10 | 0.5995 | 0.27 | **0.0097\*** | -0.13 | 0.2889 | -0.03 | 0.9299 | -0.40 | **0.0001\*** |
| Gnpda1 | -0.06 | 0.8399 | 0.11 | 0.4255 | -0.02 | 0.9155 | 0.04 | 0.9378 | -0.13 | 0.2235 |
| Gnpda2 | 0.22 | 0.5590 | 0.27 | 0.1687 | 0.31 | 0.2055 | 0.08 | 0.8864 | 0.04 | 0.8273 |
| Gpi1 | -0.10 | 0.5665 | 0.14 | 0.1191 | -0.10 | 0.4225 | 0.01 | 0.9827 | -0.24 | **0.0035\*** |
| Pfkl | -0.11 | 0.5724 | -0.37 | **0.0009\*** | -0.49 | **0.0001\*** | -0.38 | **0.0011\*** | -0.12 | 0.1415 |
| Pgam1 | -0.01 | 0.9827 | 0.03 | 0.7714 | -0.12 | 0.3132 | -0.12 | 0.5245 | -0.15 | 0.0519 |
| Pgk1 | -0.03 | 0.8623 | 0.09 | 0.2320 | -0.11 | 0.2857 | -0.08 | 0.6757 | -0.21 | **0.0041\*** |
| Pkm | -0.07 | 0.7277 | 0.12 | 0.1960 | -0.09 | 0.4372 | -0.03 | 0.9335 | -0.21 | **0.0081\*** |
| Tpi1 | -0.08 | 0.6780 | 0.25 | **0.0097\*** | -0.12 | 0.2865 | -0.05 | 0.8635 | -0.37 | **0.0001\*** |

Note: FC: fold-change; FDR: false discovery rate. Values in bold(\*) indicate the genes selected as significant from the corresponding group-wise comparisons.

Insulin treatment alone (37°C + Ins vs. 37°C) resulted in 204 differentially expressed genes (193 upregulated, 11 downregulated) (Table 1). A cluster of insulin-responsive genes (including *Egr1, Errfil1, Fos, Egr2* and *Ier2*) was highly significant (Table 2); notably, *Fos, Egr1*, and *Errfi1* were downregulated in the hyperthermia (40°C) condition compared to control (37°C), while *Arc* was significantly increased. In addition, pro-apoptotic genes Bok and Bbc3 were decreased.

The combined treatment (40°C + Ins vs. 37°C) led to 976 differentially expressed genes (694 upregulated and 282 downregulated) (Table 1). In this condition, the two most downregulated genes (*GM10532* and *Mir5114*) were no longer differentially expressed, whereas insulin-responsive genes such as *Egr1, Egr2*, and *Ier2* remained altered (Table 2). The Arc gene was notably upregulated (3.67FC). Balancing comparisons further revealed that insulin treatment in the context of hyperthermia (40°C vs. 40°C + Ins) increased the expression of numerous glycolytic pathway genes (Table 3), and that the increase in temperature in the presence of insulin (37°C + Ins vs. 40°C + Ins) mitigated some gene expression differences (Tables 2 and 3). Among all genes, *Txnip* consistently showed significant upregulation across comparisons, with a synergistic effect observed when insulin and hyperthermia were combined (40°C: +1.57 log FC; 40°C + Ins: +2.19 log FC).

### 3.3. Analysis of Shared DEGs

Analysis of shared differentially expressed genes (DEGs) across conditions revealed that the 37°C vs. 40°C + Ins comparison had the highest number of overlapping DEGs (Fig. 4a). The overlap of downregulated genes was similarly dominated first by changes in insulin status and then by temperature (Fig. 4b).

**Figure 4.**
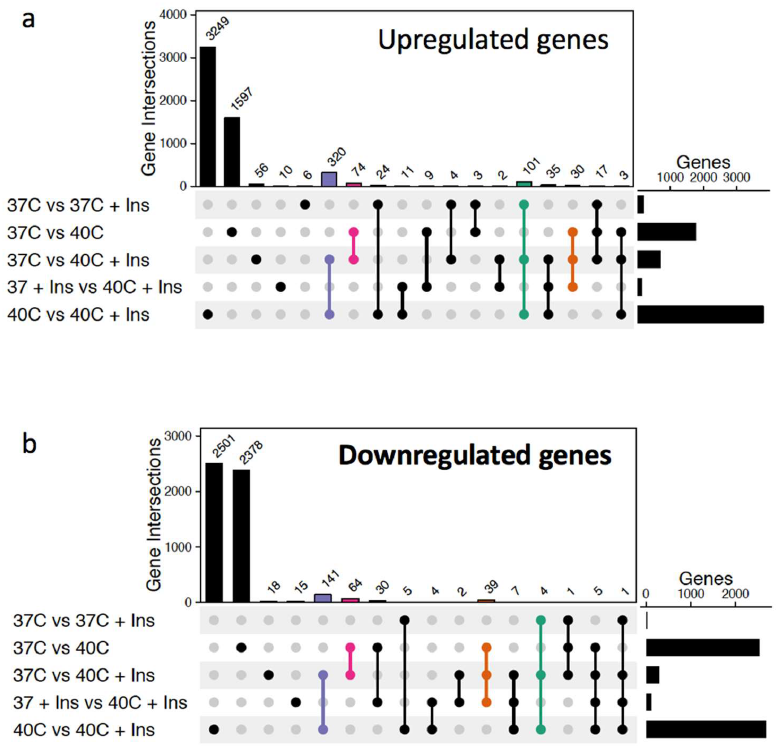
Overlap between comparisons. Upset plots showing the numbers of upregulated (**a**) and downregulated (**b**) DEGs shared across the five comparisons. The commonalities between comparisons are distinguished by changes in insulin status (purple and green) and temperature status (pink and orange).

### 3.4. Transcriptome-Wide Pathway Analysis

GO and KEGG pathway analyses of the primary comparisons identified the top 20 significantly affected networks (Fig. 5) [39]. Hyperthermia upregulated networks involved in protein synthesis and turnover (e.g., ribosome, proteasome, RNA polymerase, protein processing in the endoplasmic reticulum, spliceosome, and protein export) as well as energy metabolism (e.g., oxidative phosphorylation, thermogenesis, citrate cycle, mitophagy, pyruvate metabolism, and carbon metabolism). In contrast, networks related to cell adhesion, tight junctions, and cytoskeletal regulation were downregulated. Notably, pathways such as insulin signaling, inositol phosphate metabolism, and several proliferation-related pathways (ErbB, Hippo, and growth hormone signaling) were also down-regulated. The addition of insulin (40°C + Ins) largely abolished the temperature-induced changes, with the combined treatment clustering more closely with insulin-only profiles; for instance, networks involved in oxidative phosphorylation, thermogenesis, and ribosome function were significantly downregulated compared to hyperthermia alone.

**Figure 5.**
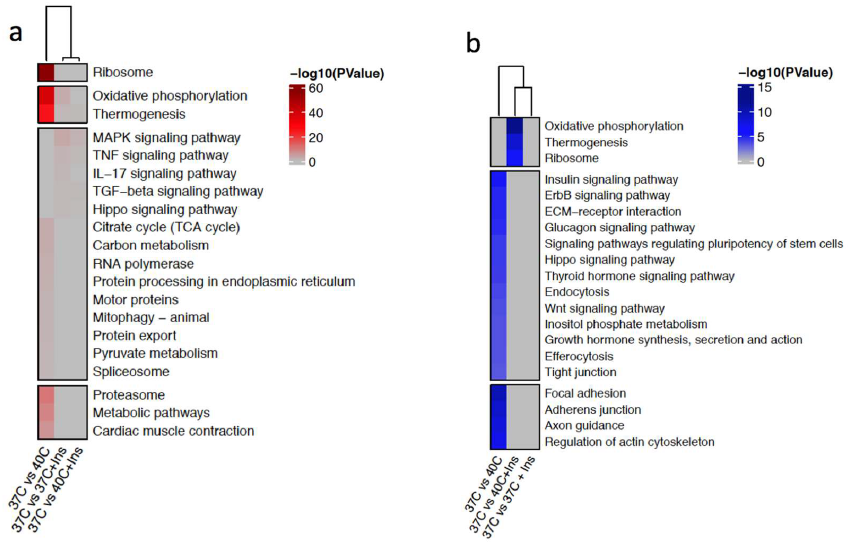
Significantly enriched KEGG Pathway Terms. Heatmaps showing the top 20 most significantly affected networks for (a) upregulated and (b) downregulated genes.

### 3.5. Targeted Gene Expression Analysis

To elucidate mechanistic insights, we focused on specific gene networks related to glucose transport, metabolism, and apoptosis.

#### 3.5.1 Glucose Import Network

Gene expression profiles were strongly influenced by insulin status, with control samples clustering closer to insulin-treated samples (Supplementary fig. S2). Under hyperthermia (40°C), genes involved in GLUT4 translocation (such as *Rab4b, Gsk3a, Rtn2*, and *Aspcr1*) were upregulated; however, this effect was largely reversed in the 40°C + Ins condition, suggesting that insulin restores these genes toward baseline levels.

#### 3.5.2 Glucose Metabolism Network

Hyperthermia alone caused distinct clustering with upregulation of genes like *Mdh1, Bad*, and *Hmgb1*, while key genes such as *Sorbs1* (important for GLUT4-mediated glucose uptake) were markedly downregulated (Supplementary fig. S3). Insulin treatment in the hyperthermia context reversed or mitigated these expression changes.

#### 3.5.3 Apoptosis Network

Consistent with the global analysis, hyperthermia induced significant upregulation of pro-apoptosis genes (*Bok, Bbc3, Trp53, Sirt2, Ptgis*, and *Gper1*) (Supplementary fig. S4). The addition of insulin reversed many of these changes, underscoring its protective role against heat-induced apoptotic signaling.

Overall, these targeted analyses indicate that insulin counterbalances the detrimental transcriptional effects of hyperthermia on glucose metabolism and apoptosis.

### 3.6 Gene Ontology Analysis

GO network analysis [37,40] was performed on DEGs (Fig. 9). Under hyperthermia (37°C vs. 40°C), genes involved in cellular respiration, mitochondrial ATP synthesis coupled to electron transport, and response to reactive oxygen species were upregulated, along with the heat response network. Networks related to autophagy and apoptosis exhibited both up- and downregulation. With the addition of insulin (37°C vs. 37°C+Ins and 37°C vs. 40°C+Ins), the apoptotic process network was further upregulated, and blood vessel morphogenesis (which was downregulated under hyperthermia alone) was altered. Overall, these analyses support that insulin mitigates many of the transcriptional changes induced by hyperthermia, particularly those related to energy metabolism and stress response.

**Figure 9.**
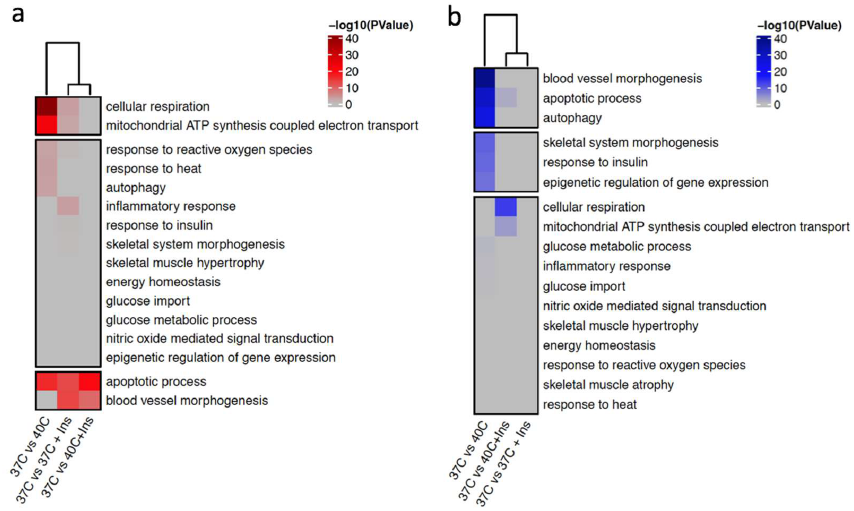
Significantly Enriched GO: Biological Function Pathway Terms. GO terms and pathway enrichment analysis based on upregulated (a) or downregulated (b) networks.

## 4. Discussion

Since their establishment by Yaffe and Saxel [10], C2C12 cells have been widely used as an in vitro model for skeletal muscle research. Our study demonstrates that acute hyperthermia induces robust transcriptional changes in C2C12 myotubes, affecting a broad range of metabolic, stress, and apoptotic pathways. Hyperthermia (40°C) significantly upregulated genes involved in glycolysis, oxidative phosphorylation, reactive oxygen species (ROS) response, and apoptosis. For instance, upregulation of genes such as *Bad, C1qtnf3, Gapdh, Tpi1, Hmgb1, Mdh2, Mdh1*, and *Trp53* suggests that cells respond to heat stress by increasing glycolytic flux and activating stress-induced apoptotic signals [41-49]. Concurrent downregulation of genes essential for normal metabolic function, including *Sorbs1* and *Atp6-ps* [50-54], further indicates that hyperthermia disrupts cellular energy regulation.

Importantly, our data reveal that the introduction of insulin during hyperthermia substantially alters this transcriptional landscape. While hyperthermia alone triggered a broad, nonspecific stress response, insulin appears to selectively modulate these effects. For example, the combined treatment (40°C + Ins) led to a significant reduction in the expression of several pro-apoptotic genes (e.g., *Bok* and *Bbc3*) compared to hyperthermia alone, and maintained or enhanced the expression of insulin-responsive genes such as *Egr1, Egr2*, and *Ier2* (Table 2). The *Arc* gene, which has been implicated in anti-apoptotic processes through its interaction with caspases [55], was also markedly upregulated when insulin was added, suggesting that insulin confers a protective effect against hyperthermia-induced cellular damage.

Targeted gene expression analyses provide further insight into the mechanisms by which insulin modulates hyperthermia responses. In the glucose import network, hyperthermia upregulated genes critical for GLUT4 translocation (e.g., *Rab4b, Gsk3a, Rtn2*, and *Aspcr1*) [56-58], yet these changes were largely reversed by insulin, returning expression profiles closer to control levels (Fig. 6). Similarly, in the glucose metabolism network, insulin treatment mitigated the hyperthermia-induced upregulation of genes such as *Mdh1, Bad*, and *Hmgb1*, and prevented the marked downregulation of key genes like *Sorbs1* (Fig. 7). These observations suggest that insulin may help restore metabolic balance under thermal stress by fine-tuning the expression of genes involved in both energy production and glucose uptake.

In addition, the global pathway analyses indicate that while hyperthermia upregulates networks associated with protein synthesis, turnover, and energy metabolism (e.g., oxidative phosphorylation, thermogenesis, and the citrate cycle), the addition of insulin shifts the overall network profile. Insulin mitigates these broad changes, downregulating pathways that were highly activated by heat stress alone (such as those related to oxidative phosphorylation and thermogenesis) and thereby potentially reducing cellular damage (Figs. 5 and 9). This shift implies a protective role for insulin, as it not only enhances glucose uptake but also helps balance the cellular energy state and reduce excessive stress responses.

In a study by Liu and Brooks (2012) [59], exposing C2C12 myotubes to mild heat stress (1 h at 40°C) led to a robust induction of mitochondrial biogenesis, as evidenced by rapid AMPK activation, increased SIRT1 and PGC-1α expression, and an elevated mitochondrial DNA copy number, effects that were most prominent 24 hours after treatment, and further enhanced with repeated exposures over five days. In contrast, our study focused on the acute transcriptional response immediately following a 30-minute hyperthermia exposure. We observed significant alterations in genes involved in glycolysis, oxidative phosphorylation, and stress responses, with insulin treatment modulating many of these hyperthermia-induced changes. Although our investigation captured early molecular events, the findings by Liu and Brooks suggest that longer recovery periods or repeated heat treatments are critical for fully manifesting the mitochondrial biogenesis program. Together, these complementary studies underscore how the timing and duration of thermal stress are key factors in determining the extent of cellular adaptation in skeletal muscle cells.

Collectively, these findings suggest that insulin exerts a dual function under hyperthermic conditions: it acts as a targeted modulator that both counterbalances detrimental transcriptional changes and supports metabolic regulation. Such a modulatory effect may be particularly relevant in physiological and pathological contexts, such as in metabolic disorders where insulin sensitivity is compromised [15-23]. These observations rely on changes in mRNA expression which do not directly reflect the expression level of the encoded proteins and their expected function, therefore further experiments to evaluate the results at the protein level are required. Also, because our conclusions are based solely on acute transcriptomic data following a 30-minute intervention, further studies incorporating time-course analyses, along with gene and protein validation experiments (qPCR and Western blot) are required to fully understand the effect of these interventions. In this pilot study, one biological replicate of the control group was excluded during quality control analysis, with only 2 samples remaining for further downstream analysis, therefore future experiments should include higher numbers of biological replicates to account for potential exclusion at the stage of quality control analysis. While our observations point to the plausible hypothesis that hyperthermia had an impact on metabolic activity and apoptosis, with increase on energy demands, more experiments are required to support this hypothesis, including assays to evaluate protein abundance, glucose metabolism, inflammation and cell survival.

Since their establishment by Yaffe and Saxel [10], C2C12 cells have been widely used as an in vitro model for skeletal muscle research. Their ease of culture, reproducibility, and well-characterized differentiation process have made them a standard model system despite their tetraploid karyotype. It is important to note that our study was conducted using C2C12 myotubes in an in vitro setting. As such, the cellular responses observed here do not fully recapitulate the complex interactions that occur in vivo, where multiple cell types—including endothelial cells, immune cells, and stromal elements—contribute to physiological outcomes such as vasodilation and systemic inflammatory responses following heat exposure. Future studies employing animal models or human tissue samples will be essential to validate these findings in the context of whole-organism physiology.

## 5. Conclusions

This study provides comprehensive insights into the acute transcriptional responses of C2C12 myotubes to hyperthermia and insulin treatment. Hyperthermia at 40°C induced significant metabolic adjustments, activated oxidative stress responses, and increased the expression of pro-apoptotic genes, reflecting the challenges of maintaining cellular homeostasis under thermal stress. Notably, the addition of insulin moderated many of these detrimental effects by enhancing glucose metabolism and reducing stress and apoptotic signaling. Although these results are preliminary and based solely on transcriptomic data, they lay the groundwork for future mechanistic studies that will incorporate protein-level and functional validations to further define the interplay between thermal stress and insulin signaling in skeletal muscle.

## Supplementary Materials

The following supporting information Figure S1, Table S1.

## Institutional Review Board Statement

Not applicable

## Informed Consent Statement

Not applicable.

## Data Availability Statement

The data are contained within the article.

---

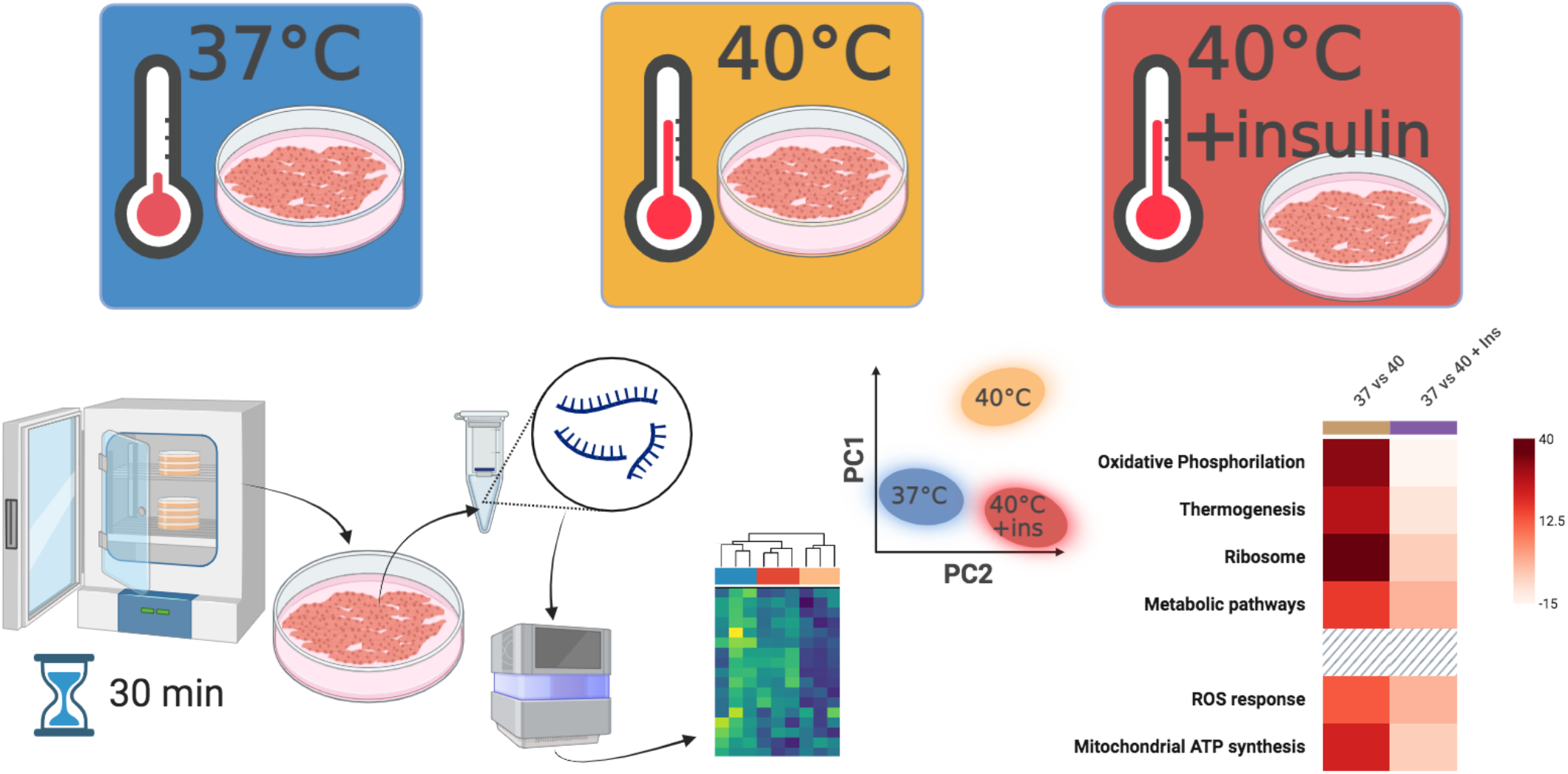

## References

1. Torday JS. Homeostasis as the Mechanism of Evolution. Biology (Basel). 2015;4(3):573–590.

2. Fulda S, Gorman AM, Hori O, Samali A. Cellular stress responses: cell survival and cell death. Int J Cell Biol. 2010;2010:214074.

3. Hildebrandt B, Wust P, Ahlers O, et al. The cellular and molecular basis of hyperthermia. Crit Rev Oncol Hematol. 2002;43(1):33–56.

4. Wang X-Y, Ostberg JR, Repasky EA. Effect of Fever-Like Whole-Body Hyperthermia on Lymphocyte Spectrin Distribution, Protein Kinase C Activity, and Uropod Formation1. The Journal of Immunology. 1999;162(6):3378–3387.

5. Sanchez-Alavez M, Tabarean IV, Osborn O, et al. Insulin causes hyperthermia by direct inhibition of warm-sensitive neurons. Diabetes. 2010;59(1):43–50.

6. Rani V, Deep G, Singh RK, Palle K, Yadav UC. Oxidative stress and metabolic disorders: Pathogenesis and therapeutic strategies. Life Sci. 2016;148:183–193.

7. Ren J, Sowers JR, Zhang Y. Metabolic Stress, Autophagy, and Cardiovascular Aging: from Pathophysiology to Therapeutics. Trends Endocrinol Metab. 2018;29(10):699–711.

8. Winn NC, Schleh MW, Garcia JN, et al. Insulin at the Intersection of Thermoregulation and Glucose Homeostasis. bioRxiv. 2023.

9. Sanchez-Alavez M, Osborn O, Tabarean IV, et al. Insulin-like growth factor 1-mediated hyperthermia involves anterior hypothalamic insulin receptors. J Biol Chem. 2011;286(17):14983–14990.

10. Yaffe D, Saxel O. Serial passaging and differentiation of myogenic cells isolated from dystrophic mouse muscle. Nature. 1977;270(5639):725–727.

11. Filigheddu N, Gnocchi VF, Coscia M, et al. Ghrelin and des-acyl ghrelin promote differentiation and fusion of C2C12 skeletal muscle cells. Mol Biol Cell. 2007;18(3):986–994.

12. Moustogiannis A, Philippou A, Zevolis E, et al. Effect of Mechanical Loading of Senescent Myoblasts on Their Myogenic Lineage Progression and Survival. Cells. 2022;11(24).

13. Huang J, Wang K, Shiflett LA, et al. Fibroblast growth factor 9 (FGF9) inhibits myogenic differentiation of C2C12 and human muscle cells. Cell Cycle. 2019;18(24):3562–3580.

14. Chai J, Xiong Q, Zhang P, Zheng R, Peng J, Jiang S. Induction of Ca2+ signal mediated apoptosis and alteration of IP3R1 and SERCA1 expression levels by stress hormone in differentiating C2C12 myoblasts. Gen Comp Endocrinol. 2010;166(2):241–249.

15. Tang J, He A, Yan H, et al. Damage to the myogenic differentiation of C2C12 cells by heat stress is associated with up-regulation of several selenoproteins. Scientific Reports. 2018;8(1):10601.

16. Lukácsi S, Munkácsy G, Győrffy B. Harnessing Hyperthermia: Molecular, Cellular, and Immunological Insights for Enhanced Anticancer Therapies. Integrative Cancer Therapies. 2024;23:15347354241242094.

17. Luo Z, Zheng K, Fan Q, Jiang X, Xiong D. Hyperthermia exposure induces apoptosis and inhibits proliferation in HCT116 cells by upregulating miR-34a and causing transcriptional activation of p53. Exp Ther Med. 2017;14(6):5379–5386.

18. Gourgou E, Aggeli IK, Beis I, Gaitanaki C. Hyperthermia-induced Hsp70 and MT20 transcriptional upregulation are mediated by p38-MAPK and JNKs in Mytilus galloprovincialis (Lamarck); a pro-survival response. J Exp Biol. 2010;213(2):347–357.

19. Lee S-H, Park S-Y, Choi CS. Insulin Resistance: From Mechanisms to Therapeutic Strategies. Diabetes Metab J. 2022;46(1):15–37.

20. Rahman MS, Hossain KS, Das S, et al. Role of Insulin in Health and Disease: An Update. Int J Mol Sci. 2021;22(12).

21. Ganesan S, Summers CM, Pearce SC, et al. Short-term heat stress altered metabolism and insulin signaling in skeletal muscle. J Anim Sci. 2018;96(1):154–167.

22. Yaribeygi H, Maleki M, Butler AE, Jamialahmadi T, Sahebkar A. Molecular mechanisms linking stress and insulin resistance. Excli j. 2022;21:317–334.

23. Kuppuswami J, Senthilkumar GP. Nutri-stress, mitochondrial dysfunction, and insulin resistance-role of heat shock proteins. Cell Stress Chaperones. 2023;28(1):35–48.

24. MoTrPAC Study Group; Lead Analysts; Amar D, Gay NR, Jean-Beltran PM, et al. Temporal dynamics of the multi-omic response to endurance exercise training. Nature. 2024;629(8010):174–183.

25. Felix Krueger FJ, Phil Ewels, Ebrahim Afyounian, Michael Weinstein, Benjamin Schuster-Boeckler, Gert Hulselmans, & sclamons. (2023). FelixKrueger/TrimGalore: v0.6.10 - add default decompression path (0.6.10). Zenodo. 10.5281/zenodo.7598955. Published 2023. Accessed.

26. Martin MCrasfh-tsrEj, 17(1), pp. 10–12. doi:10.14806/ej.17.1.200. Accessed.

27. Kim D, Pertea G, Trapnell C, Pimentel H, Kelley R, Salzberg SL. TopHat2: accurate alignment of transcrip-tomes in the presence of insertions, deletions and gene fusions. Genome Biol. 2013;14(4):R36.

28. Langmead B, Salzberg SL. Fast gapped-read alignment with Bowtie 2. Nat Methods. 2012;9(4):357–359.

29. Fgbio: Tools for Working with Genomic and High Throughput. (Fulcrum Genomics, 2024). Accessed.

30. Smith T, Heger A, Sudbery I. UMI-tools: modeling sequencing errors in Unique Molecular Identifiers to improve quantification accuracy. Genome Res. 2017;27(3):491–499.

31. Anders S, Pyl PT, Huber W. HTSeq--a Python framework to work with high-throughput sequencing data. Bioinformatics. 2015;31(2):166–169.

32. Ladwig Ww-pVxxxSB, Colorado: UCAR/NCAR. 10.5065/D6W094P1. Accessed.

33. Danecek P, Bonfield JK, Liddle J, et al. Twelve years of SAMtools and BCFtools. Gigascience. 2021;10(2).

34. Ewels P, Magnusson M, Lundin S, Kaller M. MultiQC: summarize analysis results for multiple tools and samples in a single report. Bioinformatics. 2016;32(19):3047–3048.

35. Chen Y, Chen L, Lun ATL, Baldoni PL, Smyth GK. edgeR 4.0: powerful differential analysis of sequencing data with expanded functionality and improved support for small counts and larger datasets. bioRxiv. 2024:2024.2001.2021.576131.

36. Dillies MA, Rau A, Aubert J, et al. A comprehensive evaluation of normalization methods for Illumina high-throughput RNA sequencing data analysis. Brief Bioinform. 2013;14(6):671–683.

37. Ashburner M, Ball CA, Blake JA, et al. Gene ontology: tool for the unification of biology. The Gene Ontology Consortium. Nat Genet. 2000;25(1):25–29.

38. Lund SP, Nettleton D, McCarthy DJ, Smyth GK. Detecting Differential Expression in RNA-sequence Data Using Quasi-likelihood with Shrunken Dispersion Estimates. Statistical Applications in Genetics and Molecular Biology. 2012;11(5).

39. Goto S, Bono H, Ogata H, et al. Organizing and computing metabolic pathway data in terms of binary relations. Pac Symp Biocomput. 1997:175–186.

40. Thomas PD. The Gene Ontology and the Meaning of Biological Function. Methods Mol Biol. 2017;1446:15–24.

41. Zha J, Harada H, Yang E, Jockel J, Korsmeyer SJ. Serine phosphorylation of death agonist BAD in response to survival factor results in binding to 14-3-3 not BCL-X(L). Cell. 1996;87(4):619–628.

42. Liu Z, Annarapu G, Yazdani HO, et al. Restoring glucose balance: Conditional HMGB1 knockdown mitigates hyperglycemia in a Streptozotocin induced mouse model. Heliyon. 2024;10(1):e23561.

43. Calogero S, Grassi F, Aguzzi A, et al. The lack of chromosomal protein Hmg1 does not disrupt cell growth but causes lethal hypoglycaemia in newborn mice. Nat Genet. 1999;22(3):276–280.

44. Yaribeygi H, Rashidfarrokhi F, Atkin SL, Sahebkar A. C1q/TNF-related protein-3 and glucose homeostasis. Diabetes Metab Syndr. 2019;13(3):1923–1927.

45. Jiang H, Ma N, Shang Y, et al. Triosephosphate isomerase 1 suppresses growth, migration and invasion of hepatocellular carcinoma cells. Biochem Biophys Res Commun. 2017;482(4):1048–1053.

46. White MR, Garcin ED. D-Glyceraldehyde-3-Phosphate Dehydrogenase Structure and Function. Subcell Biochem. 2017;83:413–453.

47. Jungtrakoon Thamtarana P, Marucci A, Pannone L, et al. Gain of Function of Malate Dehydrogenase 2 and Familial Hyperglycemia. J Clin Endocrinol Metab. 2022;107(3):668–684.

48. Lee SM, Kim JH, Cho EJ, Youn HD. A nucleocytoplasmic malate dehydrogenase regulates p53 transcriptional activity in response to metabolic stress. Cell Death Differ. 2009;16(5):738–748.

49. Blackburn AC, Jerry DJ. Knockout and transgenic mice of Trp53: what have we learned about p53 in breast cancer? Breast Cancer Res. 2002;4(3):101–111.

50. Baumann CA, Ribon V, Kanzaki M, et al. CAP defines a second signalling pathway required for insulin-stimulated glucose transport. Nature. 2000;407(6801):202–207.

51. Aye CC, Hammond DE, Rodriguez-Cuenca S, et al. CBL/CAP Is Essential for Mitochondria Respiration Complex I Assembly and Bioenergetics Efficiency in Muscle Cells. Int J Mol Sci. 2023;24(4).

52. Chang T-J, Wang W-C, Hsiung CA, et al. Genetic variation of SORBS1 gene is associated with glucose homeostasis and age at onset of diabetes: A SAPPHIRe Cohort Study. Scientific Reports. 2018;8(1):10574.

53. Anderson S, Bankier AT, Barrell BG, et al. Sequence and organization of the human mitochondrial genome. Nature. 1981;290(5806):457–465.

54. Nijtmans LG, Klement P, Houstek J, van den Bogert C. Assembly of mitochondrial ATP synthase in cultured human cells: implications for mitochondrial diseases. Biochim Biophys Acta. 1995;1272(3):190–198.

55. Koseki T, Inohara N, Chen S, Nunez G. ARC, an inhibitor of apoptosis expressed in skeletal muscle and heart that interacts selectively with caspases. Proc Natl Acad Sci U S A. 1998;95(9):5156–5160.

56. Eldar-Finkelman H, Krebs EG. Phosphorylation of insulin receptor substrate 1 by glycogen synthase kinase 3 impairs insulin action. Proc Natl Acad Sci U S A. 1997;94(18):9660–9664.

57. Dokken BB, Sloniger JA, Henriksen EJ. Acute selective glycogen synthase kinase-3 inhibition enhances insulin signaling in prediabetic insulin-resistant rat skeletal muscle. Am J Physiol Endocrinol Metab. 2005;288(6):E1188–1194.

58. Ikemoto T, Hosoya T, Takata K, et al. Functional role of neuroendocrine-specific protein-like 1 in membrane translocation of GLUT4. Diabetes. 2009;58(12):2802–2812.

59. Liu C-T, Brooks GA. Mild heat stress induces mitochondrial biogenesis in C2C12 myotubes. Journal of applied physiology. 2012;112(3):354–361.

